# The effect of physiological and measurement noise on the estimate of individual muscle force from indirect measurements of muscle activity

**DOI:** 10.1101/418822

**Authors:** Andrea Zonnino, Fabrizio Sergi

## Abstract

Several forward dynamics estimators have been proposed to quantify individual muscle force using non-invasive measurements of muscle activity. None of them, however, addresses the inaccuracy that arises when measurements are available only from a subset of the muscles involved in the movement under analysis. We present a novel estimator that integrates a forward dynamics estimation approach with knowledge of the optimal contraction strategy to obtain accurate estimates of individual muscle force when measurements of muscle activity are not available for all muscles. A following *in-silico* characterization showed that when trying to estimate forces form the forearm muscles acting around the wrist joint, our novel estimator is able to decrease the mean estimation bias by about 25% of the true value of muscle force. With a sensitivity analysis, we show that the model-based estimator is robust against physiological variability in muscle co-contraction strategy.

## 1. Introduction

Quantification of individual muscle force applied during tasks that require coordinated muscle co-activation would provide considerable insights in neuromuscular physiology, and enable accurate diagnosis and management of neuromotor disorders. However, direct measurement of individual muscle force is not possible *in-vivo* since it requires invasive procedures to place force sensing elements in series to the musculo-tendon units (Ravary et al. 2004; Fleming and Beynnon 2004; Dennerlein et al. 1998; Dennerlein 2005). Consequently, quantifying individual muscle force during complex tasks still represents one of the biggest challenges in biomechanics (Herzog 2017).

Alternatively, non-invasive approaches have been proposed to estimate individual muscle force using estimation techniques (Erdemir et al. 2007). These methods rely on the evidence that muscle forces produce joint movement and torque. However, since neither joint position nor joint torque provide information at the individual muscle level, these measurements need to be augmented via estimators to quantify the individual muscle forces.

Current estimation algorithms can be grouped in two macro-categories: inverse and forward dynamics approaches. Inverse dynamics estimators use either the measured joint torque or joint angle to estimate the contribution of different muscles to the measured movement (Anderson and Pandy 2001; Thelen et al. 2003; Tsirakos et al. 1997). Cost-function minimization is usually employed to address muscle redundancy (Winter 2005); unfortunately, the assumption of a cost function that is generalizable across tasks and subjects does not always hold, with the estimated forces highly sensitive to the selection of such function (Collins 1995; Glitsch and Baumann 1997).

Forward dynamics estimators (Sartori et al. 2012; Buchanan et al. 2005; Winby et al. 2009; Lloyd and Besier 2003), on the contrary, make no assumption on the strategy adopted to solve the muscle redundancy. They typically require knowledge of the limb’s geometry, measurements of joint torque, and measurements of muscle activity, typically obtained with surface electromyography (sEMG). However, since sEMG can only measure the activity of superficial muscles, current estimation approaches neglect the contribution from non-superficial muscles to the joint torque. This approximation might be appropriate when estimating forces from the lower limbs muscles (Sartori et al. 2012), however it is likely to result in inaccurate estimates when applied to complex body segments that have many muscles arranged in different layers, such as the forearm muscles that control hand and wrist movements.

In this paper, we present a novel estimator that integrates a forward dynamics estimation approach with a neuromuscular model of muscle contraction, used to estimate the effect of unmeasured muscle activity on joint torque. We hypothesized that the novel estimator would be able to return more accurate estimates of muscle force compared to a standard musculoskeletal estimator, when both are used to estimate forces of the forearm muscles during isometric tasks involving the wrist. Since the true value of the muscle force is not measurable through *in-vivo* procedures, it would be impossible to quantify accuracy of any muscle force estimator using purely experimental protocols. As such, we chose to simulate measurements using a musculoskeletal model, which allowed to run simulations to quantify the deviation between the estimated and the assumed true values of muscle forces. We thus developed a computational framework based on a realistic musculoskeletal model (Saul et al. 2014) that simulates muscle forces and virtual measurements of muscle activity during isometric contractions. To test our hypothesis, we simulated calibration experiments aimed at estimating individual muscle force from a set of measurements of muscle activity under a set of experimental conditions, when realistic conditions of measurement noise and physiological variability are included.

## 2. Materials and Methods

### 2.1. Model

The computational framework presented in this paper is based on the upper extremity musculoskeletal model presented by Saul et. al (Saul et al. 2014). Only a subset of the objects in the model are used for this analysis; specifically, the upper arm is considered grounded, with motion allowed only for the forearm and hand. As such, the reduced model has four degrees of freedom (DOFs): elbow flexion/extension (eFE), pronation/supination (PS), wrist flexion/extension (FE), and wrist radio/ulnar deviation (RUD). A total of *m_tot_* = 24 muscle segments are included (Tab. 1); however, for some of the analyses presented, a further reduced set of muscles will be considered.

**Figure 1.**
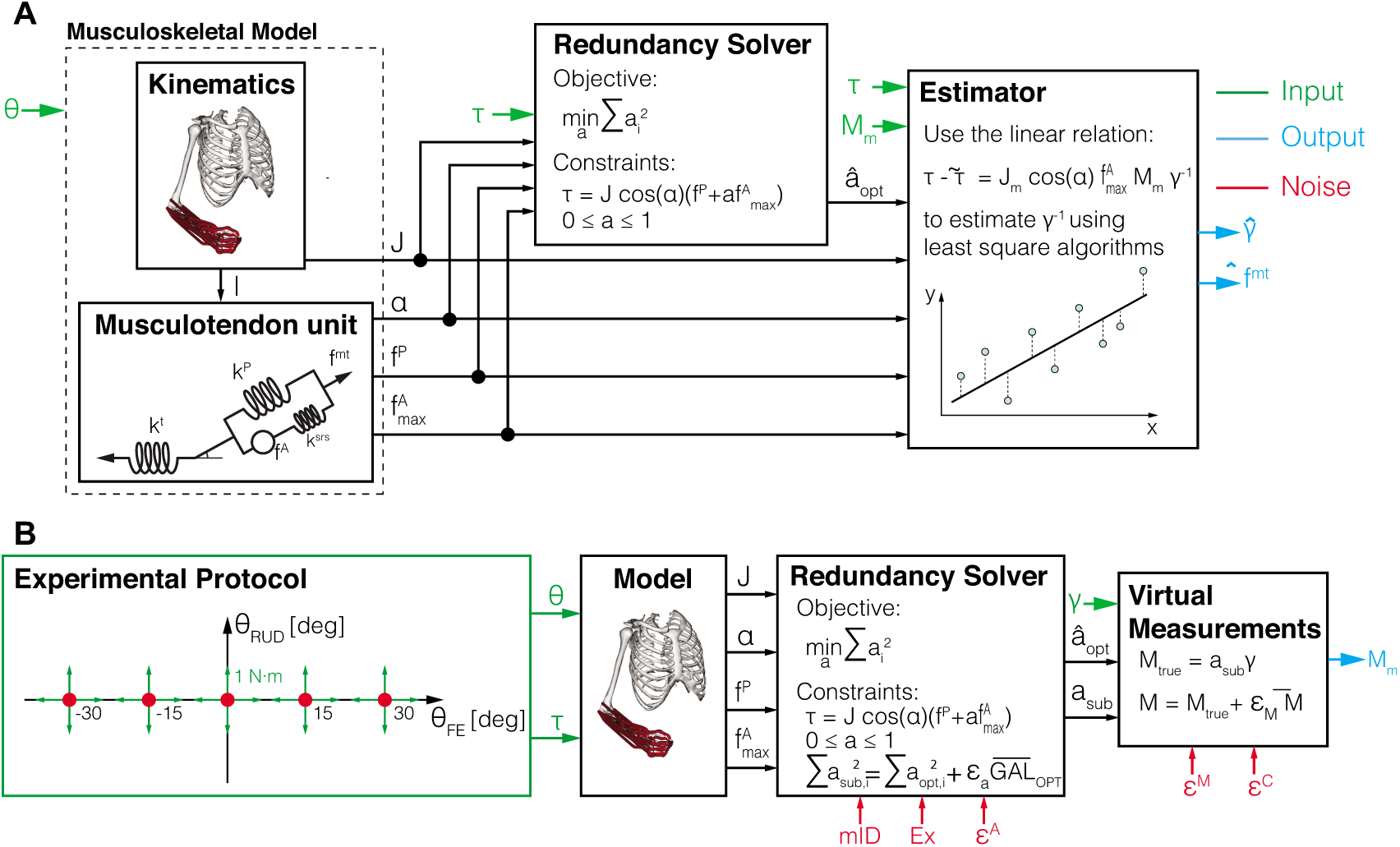
The block diagram (A) shows the schematic structure of the proposed estimator. The block diagram (B) shows the the different steps needed to generate the virtual measurements used to validate the proposed estimator. In the figure the parameters *ϵ*^*A*^, *ϵ*^*C*^ *ϵ* ^*M*^ represents the level of physiological variability, cross-talk, and measurements error, respectively; mID is a vector that contains the set of measurable muscles, and Ex is a boolean variable that indicate if the contribution of the finger muscles is negligible (0) or not (1).

Each muscle segment included in the MSM is modeled as a Hill-type musculotendon (MT) unit in isometric conditions (Hill 1938; Millard et al. 2013). For given values of muscle activation *a* and muscle length *l*_*m*_, the musculo-tendon force *f*_*MT*_ of each segment is determined by:

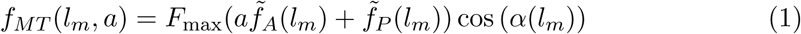

where *F*_max_ is the maximum isometric muscle force, *α* is the muscle pennation angle, and 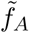 and 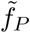 are the active and passive force multipliers, respectively (Millard et al. 2013). Assuming the tendon to be inextensible (Seth et al. 2011), the length of each muscle is directly related to the posture of the wrist joint and independent from muscle activation. As such, in a given posture the MT force is linearly proportional to muscle activity.

To simulate indirect measurements of muscle activity, a linear dependence is then assumed between the value of activation *a* and the measurement *M:*

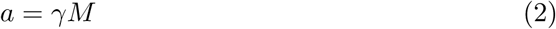

where γ is a muscle-specific scaling coefficient that needs to be derived via a proper calibration procedure to estimate activation and force from the measured values of *M.*

### 2.2. Muscle Force Estimation

In general, the relationship between muscle force and joint torque is provided by:

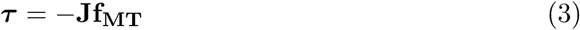

As such, combining eq. (1) - (3), and using the bold notation to refer to vector or matrix quantities, a linear relationship can be derived between τ and ***M***:

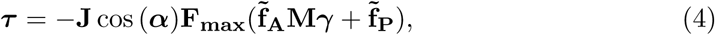

where **J** is the muscular Jacobian whose component *r_ij_* represents the moment arm of the muscle *j* with respect of the joint angle *i.* **F**_**max**_, α, and 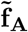 are diagonal matrices that contain the respective scalar parameters for each muscle. γ and 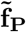 are vectors that contain the muscle-specific scaling coefficient and passive force for each muscle, respectively.

From eq. (4), it can be seen that there is a linear relationship between τ and M. As such, by measuring joint torque, measuring the activation of all muscles producing torque about a given joint, and accounting for the passive force of all such muscles, all elements in the vector γ can be estimated with a proper calibration procedure.

#### 2.2.1. Musculoskeletal estimator (MSK)

While the general description reported above is theoretically correct, the assumptions included therein are almost never achievable in practice. In fact, standard non-invasive muscle activation measurement techniques, such as sEMG, can only provide a measurement of the activation of superficial muscles. Usually, activation of *m < m*_*tot*_ muscles is available using sEMG. A standard approach to calibration, in these cases, neglects the contribution of unmeasured muscles and of passive forces, simplifying eq. (4) as

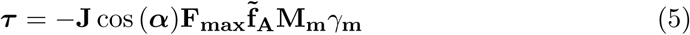

where **M**_m_ and γ_*m*_ are the subsets of M and γ that pertain to the measurable muscles.

Since eq. (5) is a linear equation of the form **y** = **X***β*, with proper experimental design, it is possible to define a set of n conditions, defined as a set of joint torques applied at different postures in isometric conditions, such that the resulting experimental matrix 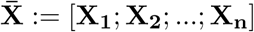 is of full rank. When this condition is satisfied, it is possible to estimate the vector *β* := γ_*m*_ using a standard least squares fit given measurements of 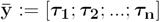.

#### 2.2.2. Neuromusculoskeletal estimator (NMSK)

We propose a novel estimation algorithm capable of dealing with cases where the approximations made within the MSK estimator do not allow sufficient accuracy, due to unmeasured muscle activations. Instead of neglecting the contribution of unmeasured muscles and of passive muscle forces, our new estimator integrates quantities available through measurements with parameters extracted from a musculoskeletal model to improve the accuracy of the estimation of vector γ_m_. The estimator extracts the contribution of unmeasured muscle forces using a neural model that provides a solution for the muscle redundancy problem. As such, we refer to our estimator as neuromusculoskeletal estimator (NMSK).

A schematic of the NMSK estimator is shown in Fig. 1. For a generic condition, defined by a specific value of joint angle and torque, static optimization is used to obtain the optimal muscle activation vector, 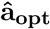, that minimizes the Global Activation Level 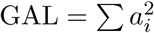 (Seth et al. 2011). The subset 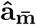 of 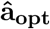 that correspond to unmeasured muscles are then included in eq. (4), together with the model-estimated passive muscle force:

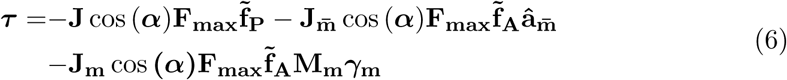

where the subcomponents of the Jacobian corresponding to measured and unmeasured muscles, **J**_**m**_ and 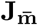 respectively, have been decoupled.

The parameters in the first two terms are obtained from the MSM and are constant for a given condition. Thus, if we define the quantity 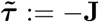 cos (α) 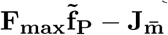 cos (α) 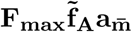, eq. (6) simplifies to:

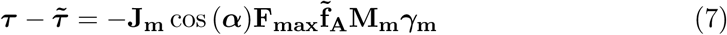

which is a linear equation of the form **y** = **X***β*, which is in the same form as eq. (5). As such, the vector Y_m_ can be estimated based on measurements of ***τ*** and **M**_**m**_ using the same experimental design described for the MSK estimator.

### 2.3. Model-based characterization of the estimators

To investigate the accuracy of our proposed estimator compared to the standard estimation methods, we simulated virtual calibration experiments, where the quantities **M** and **τ** are measured for a given experimental protocol, composed of *N* experimental conditions. Each experimental condition is defined as an isometric contraction, applied at a given joint posture under the application of a given torque. For each condition, the MSM produces simulated values of muscles activity for all muscles using its standard solver (Seth et al. 2011), based on the minimization of the global activation level 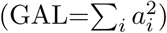. For a given experimental design, the accuracy of the estimate will largely depend on the validity of the estimator assumptions and on the precision of the measurements. We have, then, introduced several sources of non-ideality in our simulations to quantify the effect of physiological variability and measurement error under different experimental conditions on the accuracy of the estimators.

**Figure 2.**
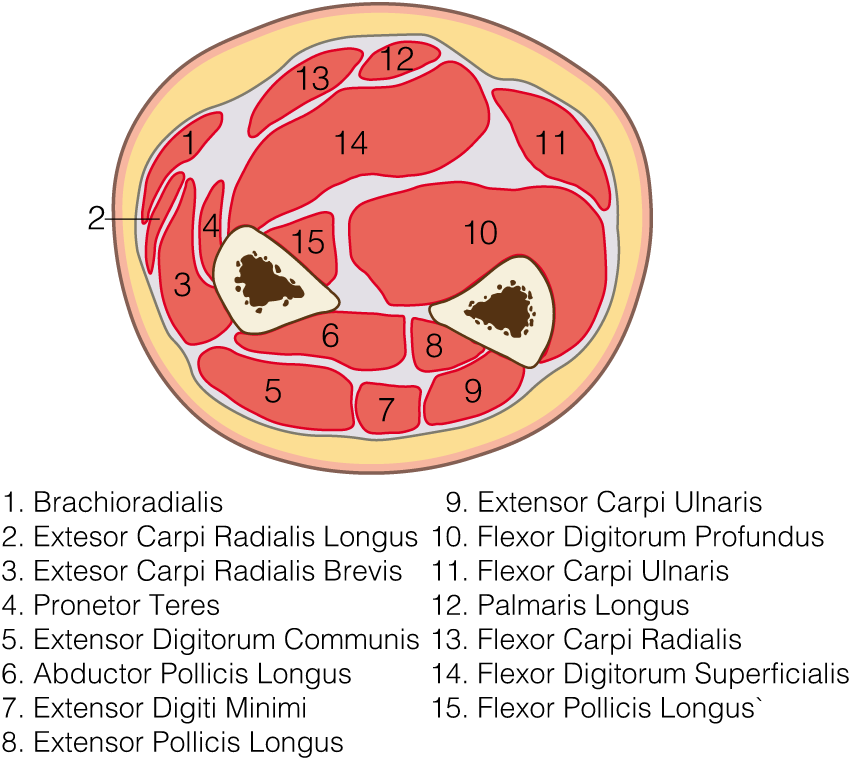
Schematic of the transverse section of the middle forearm

#### 2.3.1. Experimental conditions

An important factor that is expected to affect the accuracy of the estimation relates to the way the arm is constrained during the isometric contractions. In the specific case of the forearm, finger muscles have moment arm about the wrist, but they are not always accessible using sEMG (see Fig. 2 and Tab. 1). As such, unmeasured activity from finger muscles could have an effect on calibration procedures that relate measured activation to wrist joint torque. Activity from finger muscles can be mediated by proper experimental design: if the fingers are unconstrained and subjects are instructed to apply wrist torque without moving their fingers, we expect that finger muscle activations will be much lower than if the fingers and palm are all constrainedin a single posture (e.g. using a rigid support for the hand). In our framework, we implemented the possibility to include or neglect the contribution of finger muscles when generating virtual measurements for models that include a different total number of muscles *mtot* (factor Ex).

#### 2.3.2. Measurement properties

The quality of estimation crucially depends on how measurements of muscle activity are obtained, which is defined in terms of which muscles are measured (Me_1_), and how much noise is included in these measurements (Me_2_ and Me_3_).

We considered the effect of two cascaded sources of noise: one related to the crosstalk between each muscle and those surrounding it (Me_2_), and the other related to measurement error at the individual muscle level (Me_3_). To define levels of factor Me_2_, we assumed that the measurement of the activation for a given muscle is affected by the activity of surrounding muscles as described by the equation:

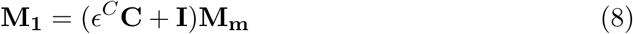

where ϵ^*C*^ is a gain factor, **I** is the identity matrix, 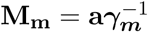 is the measurement corresponding to the true value of activation **a**, and **C** is a cross-correlation matrix that represents how much the measurement *M*_1_ of one muscle is affected by the activation of all other muscles in the limb. For proper scaling of within-and across-muscle errors, the sum of the elements for each row have unitary magnitude, with zeros in the diagonal. We heuristically defined the values of the off-diagonal elements of the cross-correlation matrix based on proximity and relative size of the pairs of muscles (Tab. 2).

To define levels of factor Me_3_, we considered that the final measurement *M*_*2*_ is affected by measurement noise, as defined by

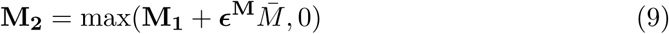

where **M**_**1**_ is the vector obtained using eq. (8) and 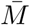 is the average value of the measurements **M**, simulated in the optimal control strategy, across muscles and conditions.

**Table 1.** Muscle groups in the model

| Group | Muscle Name | Abbreviation |
| --- | --- | --- |
| **Wrist** | Extensor Carpi Radialis Brevis | ECRB |
|  | Extensor Carpi Radialis Longus | ECRL |
|  | Extensor Carpi Ulnaris | ECU |
|  | Flexor Carpi Radialis | FCR |
|  | Flexor Carpi Ulnaris | FCU |
| **Palm** | Palmaris Longus | PL |
| **Thumb** | Extensor Pollicis Brevis | EPB |
|  | Extensor Pollicis Longus | EPL |
|  | Abductor Pollicis Longus | APL |
|  | Flexor Pollicis Longus | FPL |
| **Finger** | Extensor Digitorum Communis* | EDC |
|  | Extensor Indicis Proprius | EIP |
|  | Extensor Digiti Minimi | EDM |
|  | Flexor Digitorum Superficialis* | FDS |
|  | Flexor Digitorum Profundus* | FDP |

\* muscle composed of different segments, one per each finger: Index (I), Middle (M), Ring (R), Little (L). Since the model does not describe individual finger motion, a single muscle including all these segments is considered in this work. 2

**Table 2.** Definition of the matrix C

|  | ECRB | ECRL | ECU | FCR | FCU | PL | EPL | FPL | APL | FDS | FDP | EDC | EIP | EDM |
| --- | --- | --- | --- | --- | --- | --- | --- | --- | --- | --- | --- | --- | --- | --- |
| ECRB | 0 | 0.25 | 0 | 0 | 0 | 0 | 0 | 0.1 | 0.15 | 0.1 | 0 | 0.4 | 0 | 0 |
| ECRL | 0.5 | 0 | 0 | 0 | 0 | 0 | 0 | 0.1 | 0 | 0 | 0 | 0.4 | 0 | 0 |
| ECU | 0 | 0 | 0 | 0 | 0 | 0 | 0.2 | 0 | 0.1 | 0 | 0.5 | 0 | 0 | 0.2 |
| FCR | 0.1 | 0.05 | 0 | 0 | 0 | 0.3 | 0 | 0 | 0 | 0.55 | 0 | 0 | 0 | 0 |
| FCU | 0 | 0 | 0 | 0 | 0 | 0 | 0 | 0 | 0 | 0.5 | 0.5 | 0 | 0 | 0 |
| PL | 0 | 0 | 0 | 0.3 | 0.15 | 0 | 0 | 0 | 0 | 0.55 | 0 | 0 | 0 | 0 |
| EPL | 0 | 0 | 0.2 | 0 | 0 | 0 | 0 | 0 | 0.35 | 0 | 0.35 | 0 | 0 | 0.1 |
| FPL | 0 | 0 | 0 | 0 | 0 | 0 | 0 | 0 | 0.1 | 0.45 | 0.45 | 0 | 0 | 0 |
| APL | 0.15 | 0 | 0 | 0 | 0.05 |  | 0.15 | 0.15 | 0 | 0 | 0 | 0.4 | 0 | 0.1 |
| FDS | 0.05 | 0.05 | 0 | 0.1 | 0.2 | 0.1 | 0 | 0.1 | 0 | 0 | 0.4 | 0 | 0 | 0 |
| FDP | 0 | 0 | 0.1 | 0 | 0.25 | 0 | 0.1 | 0 | 0.1 | 0.45 | 0 | 0 | 0 | 0 |
| EDC | 0.3 | 0 | 0.05 | 0 | 0 | 0 | 0.05 | 0 | 0.4 | 0 | 0 | 0 | 0 | 0.2 |
| EIP | 0 | 0 | 0.2 | 0 | 0 | 0 | 0 | 0 | 0.1 | 0 | 0 | 0.5 | 0 | 0.2 |
| EDM | 0 | 0 | 0.2 | 0 | 0 | 0 | 0.2 | 0 | 0.3 | 0 | 0 | 0.3 | 0 | 0 |

**Table 3.** Results of the general lineal model _tting

|  | Estimate | SE | tStats | p |
| --- | --- | --- | --- | --- |
| Intercept | 0.0002 | 0.0052 | 0.0452 | 0.9639 |
| E | 0 | 0.0073 | 0 | 1 |
| Ex | 0.0033 | 0.0077 | 0.4351 | 0.6635 |
| Me<sub>1</sub> | 0.0297 | 0.0073 | 4.0318 | <0.001 |
| Ph | 0.0014 | 0.0142 | 0.1026 | 0.9183 |
| Me<sub>2</sub> | 0.0002 | 0.0141 | 0.0155 | 0.9876 |
| Me<sub>3</sub> | 0.1052 | 0.0142 | 7.3982 | <0.001 |
| E · Me<sub>1</sub> | -0.0236 | 0.0104 | -2.2718 | 0.0233 |
| E · Ex | 0 | 0.0109 | 0 | 1 |
| E · Ph | 0 | 0.0209 | 0.0200 | 1 |
| E · Me<sub>2</sub> | 0 | 0.0200 | 0 | 1 |
| E · Me<sub>3</sub> | 0 | 0.0201 | 0 | 1 |
| Ex · Me<sub>1</sub> | 0.2772 | 0.0106 | 25.9370 | <0.001 |
| Ex · Ph | 0.0680 | 0.0209 | 3.2462 | 0.0012 |
| Ex · Me<sub>2</sub> | 0.2723 | 0.0210 | 12.9150 | <0.001 |
| Ex · Me<sub>3</sub> | 0.7847 | 0.0229 | 34.1810 | <0.001 |
| Me<sub>1</sub> · Ph | 0.0445 | 0.0200 | 2.2186 | 0.0267 |
| Me<sub>1</sub> · Me<sub>2</sub> | -0.0014 | 0.0200 | -0.0696 | 0.9444 |
| Me<sub>1</sub> · Me<sub>3</sub> | -0.0656 | 0.0201 | -3.2638 | 0.0011 |
| E · Ex · Me<sub>1</sub> | -0.2785 | 0.0151 | -18.433 | <0.001 |
| E · Ex · Ph | 0 | 0.0296 | 0 | 1 |
| E · Ex · Me<sub>2</sub> | 0 | 0.0298 | 0 | 1 |
| E · Ex · Me<sub>3</sub> | 0 | 0.0324 | 0 | 1 |
| E · Me<sub>1</sub> · Ph | 0.0181 | 0.0284 | 0.6383 | 0.5234 |
| E · Me<sub>1</sub> · Me<sub>2</sub> | 0.0002 | 0.0282 | 0.0061 | 0.9950 |
| E · Me<sub>1</sub> · Me<sub>3</sub> | 0.0158 | 0.0284 | 0.5581 | 0.5769 |
| Ex · Me<sub>1</sub> · Ph | 0.0053 | 0.0290 | 0.1833 | 0.8545 |
| Ex · Me<sub>1</sub> · Me<sub>2</sub> | -0.0644 | 0.0292 | -2.2044 | 0.0277 |
| Ex · Me<sub>1</sub> · Me<sub>3</sub> | -0.8526 | 0.0305 | -27.9280 | <0.001 |
| E · Ex · Me<sub>1</sub> · Ph | -0.0399 | 0.0411 | -0.9692 | 0.8545 |
| E · Ex · Me<sub>1</sub> · Me<sub>2</sub> | 0.0219 | 0.0412 | 0.5306 | 0.5957 |
| E · Ex · Me<sub>1</sub> · Me<sub>3</sub> | 0.3096 | 0.0431 | 7.1730 | <0.001 |

The gain factor **ε**^***M***^ is a diagonal matrix.

#### 2.3.3. Physiological variability

Physiological variability (Ph) accounts for the variability associated with the selection of the contraction strategy for tasks that require coordinated muscles activation. We modeled this variability assuming that, when solving for activation values compatible with the target joint torque, the virtual subject selects a suboptimal solution (different from the optimal one where, whose ‘suboptimality level’ is defined in term of percentage of additional GAL used to complete the isometric task:

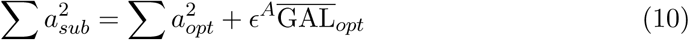

where 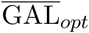 is the average GAL_*opt*_ across the different conditions, and *ε*^*A*^ a gain factor.

### 2.4. Sensitivity analysis

We conducted virtual calibration experiments where muscle activity is simulated under different levels of the previously described factors to assess the robustness of the two estimators to changes in experimental conditions, measurement properties, and physiological variability.

Factor levels are defined as follows. We considered two levels of factor Ex, i.e. fingers unconstrained/constrained. In the unconstrained mode, we assumed negligible the activation of finger muscles and their passive forces. In the constrained mode, the fullmodel (forearm, hand, fingers) was considered. Two different levels for factor Me_1_ were considered, the first level corresponding to the case where measurements are available only from the five main wrist muscles (MeasWrist), i.e. ECRL, ECRB, ECU, FCR, FCU, and the second level corresponding to the case where measurements are available from all muscles included in the model (MeasAll).

Values of factors representing a continuous source of noise/variability were selected from uniform distributions, 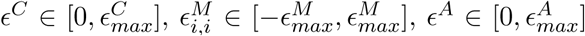, with a different value selected for each condition. For all three factors, five levels of 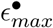 were used 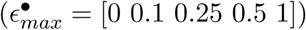.

This analysis results in a full factorial experimental design (5×5×5×2×2) composed of 500 combinations. For each combination, we simulated *N* = 25 virtual experiments, each composed of 25 isometric contractions, applied in five different wrist postures ([*θ*_*FE*_, *θ*_*RUD*_] ∈ {[—30, 0], [—15,0], [0,0], [15,0], [30, 0]}), when constant torque was applied along each of the four cardinal directions (pure wrist FE and wrist RUD torque in both directions), with unitary magnitude, followed by a rest condition (zero joint torque). For each contraction, we simulated measurements of muscle activity by assuming *γ_m_* = 10 for all measured muscles.

For each experiment, we estimated 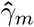 using both MSK and NMSK, and quantified the accuracy of each estimator as the bias 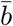, defined as the percent error in estimating γ _m_ for the five main wrist muscles, averaged across muscles.

#### 2.4.1. Statistical model fit

To establish the relationship between the factors and the outcome metric 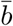, we fit a General Linear Model (GLM) to the simulated data. Ph, Me_2_, and Me_3_ were modeled as ordinal variables characterized by the mean value of |ϵ^•^ | across the 25 experimental conditions. Ex and Me_1_ are instead modeled as categorical variables. Specifically, Ex assumes the value of 0 if the fingers joint are constrained and 1 if they are unconstrained, while Me_1_ assumes value of 0 if the measurements are available from all muscles and 1 if they are available only from the five main wrist muscles. Moreover, a third categorical variable E has been used to include the factor ‘estimator’ in the GLM; we assigned the value of 0 to MSK and the value of 1 to NMSK.

To quantify the relationship between estimation bias 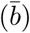, measurement error (Me_2_, Me_3_), and physiological variability (Ph), under all possible experimental and measurement conditions encoded by categorical variables (E, Ex, Me_1_), we created a GLM that included all interactions between the three categorical variables, as well as the interactions between all combinations of categorical variables and Me_2_, Me_3_, Ph, separately. This resulted in a model with 32 factors, which included 3 four-way interaction terms, 10 three-way interaction terms, 12 two-way interaction terms, in addition to the linear contribution of the 6 factors and the intercept. Statistically significant association between the metric 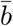 and each term in the GLM was considered for a type-I error rate smaller than 0.05.

To allow for visualization and interpretation of effects, for each significant term, we defined residual bias as the difference between the simulated bias 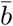, and the bias *b*_*other*_ calculated as the bias explained by all terms that do not include any factor in the significant term (e.g. for the term E-Me_1_, we would include in *b*_*other*_ the components estimated by the model that do not include factors E and Me_1_). This operation was conducted assuming that all other factors in the model were at their average level. This allowed to obtain distributions of residual bias for each combination of factors included in the significant term (2 for a main effect, 4 for a two-way interaction, etc.), which could be visually checked for model fit by comparing the distribution of 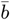 with the corresponding model values, and to identify which combination of factors contributes to the significance of the term.

**Figure 3.**
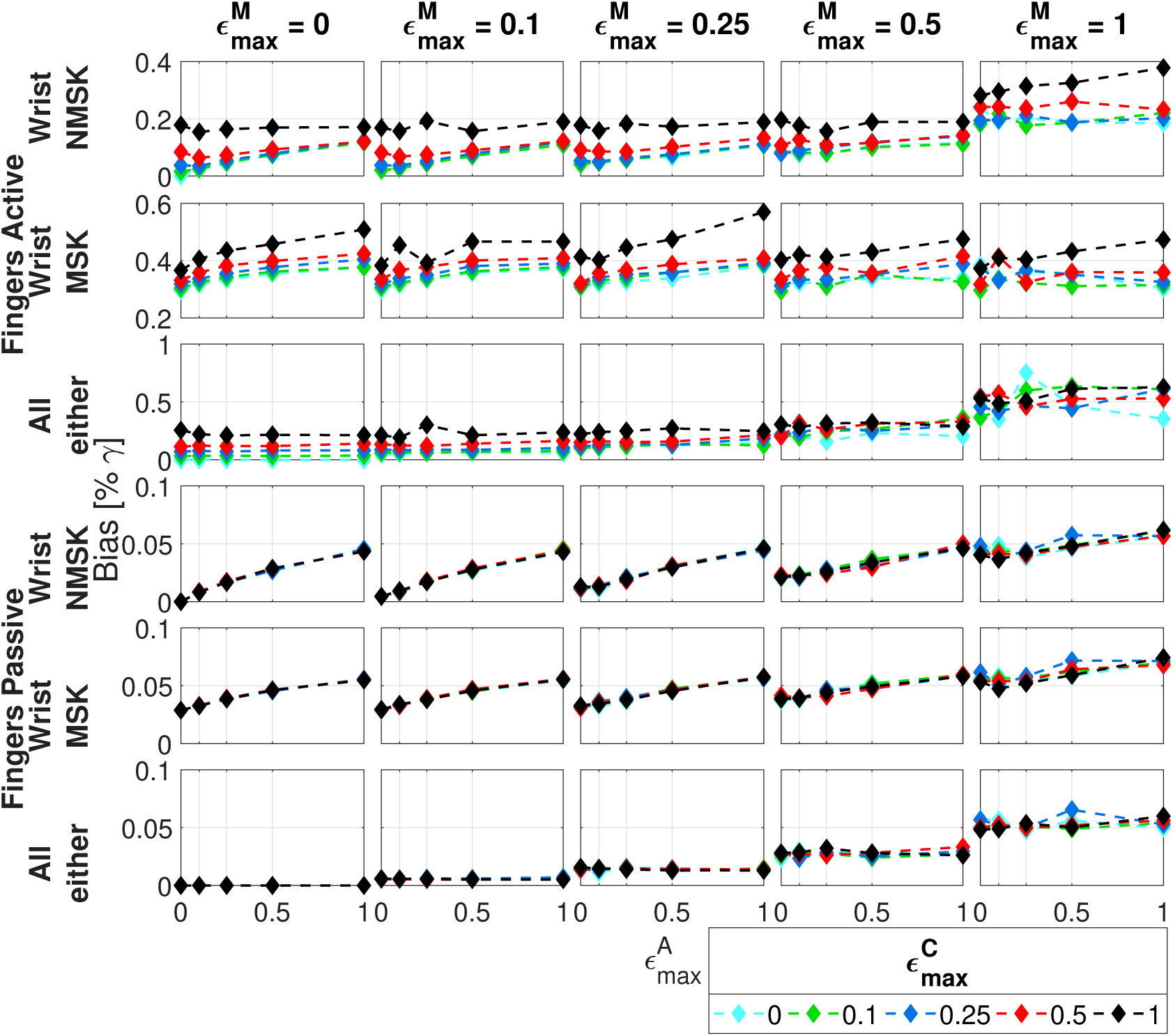
Values of metric 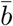 obtained for all levels of all factors. The top three rows include results obtained in presence of finger muscles (Ex=1), while the bottom three rows report the results obtained in the absence of finger muscles (Ex=0). The first two rows of each of subgroup report bias obtained when measurements are available from only wrist muscles (Me_1_ = 1), while the third row reports results obtained when measurements are available from all muscles (Me_1_ = 0). Rows one and four include estimation results obtained with the NMSK estimator (E=1), while rows two and five are obtained with the MSK estimator (E=0). Rows three and six are obtained with either estimator since no difference is expected when measurements are available from all muscles. Columns encode levels of factor Me_3_ (measurement error), colors encode levels of factor Me_2_ (cross-talk), while the x-axis of each plot encodes levels of factor Ph (physiological variability).

## 3. Results

The results of the sensitivity analysis are graphically summarized in Fig. 3, where each dot represents the average value of the metric 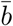 obtained for each set of factors in the 25 different repetitions. The GLM fit the simulated data with an *R*^*2*^ = 0.96. The coefficients estimated for all model terms, along with the standard error (SE), the test statistics (tStats) and the level of significance (two-sided tests) are reported in Tab. 3.

Model fit identified a significant main effect of only two of the six factors, Me_1_ and Me_3_ (p <0.001 for both) (Fig. 4). Specifically, the bias explained by only factor Me_1_, when all other factors are in their average conditions, was significantly smaller in the MeasAll condition than in the MeasWrist condition (mean ± 95% confidence interval - 0.11±0.01 MeasAll, vs. 0.14±0.01 MeasWrist). For Me_3_, each unit increase in measurement error *eu* resulted in an increase in residual bias of 0.29 when all other factors were in their average conditions.

**Figure 4.**
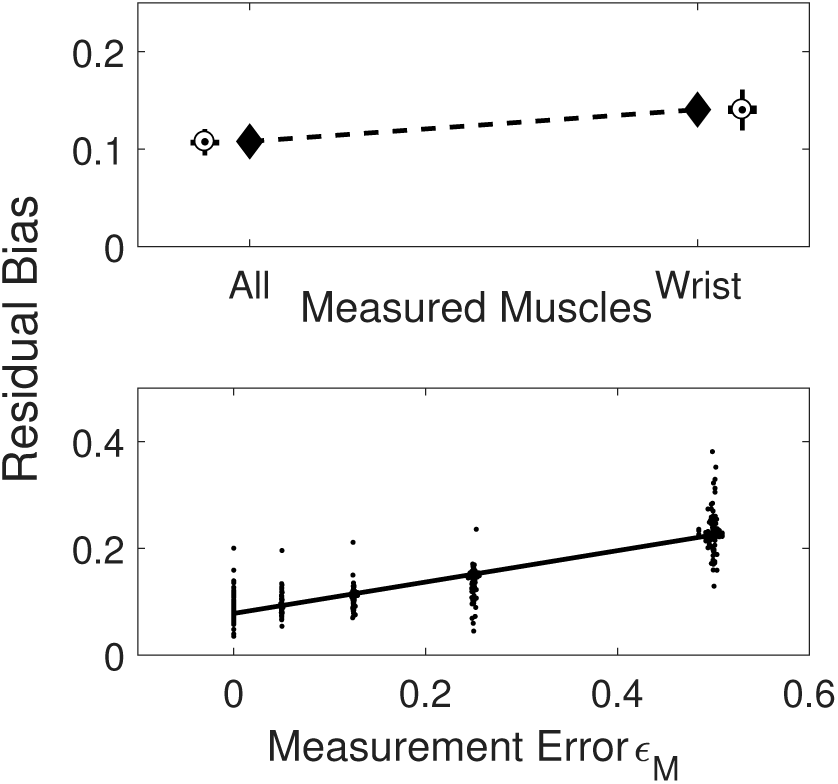
Graphical representation of the linear contribution of the two statistically significant predictors. In the top row the diamonds represents the model estimates, with the simulated data reported in the error bar. In the bottom row the line represents the model estimate while the points are the simulated data. All plots are obtained in the average condition of noise for the unreported variables

Seven two-way interaction terms had a statistically significant effect, i.e. E·Me_1_ (p = 0.0233), Ex·Me_1_ (p <0.001), Ex·Ph (p = 0.0012), Ex·Me_2_ (p <0.001), Ex·Me_3_ (p <0.001), Me_1_·Ph (p = 0.0267), Me_1_·Me_3_ (p = 0.0011). Further analysis of the effects of the two-way interactions is excluded from this section for the sake of space. All significant effects are broken down in the interaction plots shown in Fig. 5, when all other factors are in their average condition.

Model fit identified as statistically significant three three-way interaction terms, i.e. E·Ex·Me_1_ (p <0.001), Ex·Me_1_·Me_2_ (p = 0.0277), Ex·Me_1_·Me_3_ (p <0.001) (Fig. 6), and one four-way interaction term, E·Ex·Me_1_·Me_3_ (p <0.001) (Fig. 7). For the interaction between E, Ex, and Me_1_ (Fig. 6, top row), the NMSK estimator introduced a statistically significant decrease in estimation bias over the MSK estimator in the MeasWrist condition (0.05±0.01 MSK vs. 0.03±0.01 NMSK, fingers passive; and 0.36±0.02 MSK vs. 0.12±0.02 NMSK, fingers active), but the effect of the estimator was greater when fingers were active than when fingers were passive (change in bias: 0.02 ± 0.01 fingers passive, vs. 0.24±0.03 fingers active). As expected, no difference between the two estimators was observed in the MeasAll condition; in that condition, bias was greater when fingers were active than when they were passive (0.2±0.02, fingers active vs. 0.02±0.02, fingers passive).

For the interaction between Me_1_, Ex, and Me_2_ (Fig. 6, middle row), cross-talk introduced no significant effect on bias when fingers were passive (slope: 0± 0.1 in both cases), but bias significantly increased in the MeasWrist condition (bias: 0.02±0.01 MeasWrist vs. 0.03±0.01 MeasAll). Instead, when fingers were active, a significant effect of cross-talk was measured in both the MeasAll and the MeasWrist condition (slope: 0.22±0.08 for MeasWrist vs 0.27±0.09 for MeasAll). However, no significant difference between the two slopes was measured. In general, regardless of the value of cross-talk, in the finger active condition, bias in the MeasAll condition was always larger than bias in the MeasWrist condition (bias: 0.19±0.01 for MeasAll vs. 0.24±0.01 MeasWrist), as seen in Fig. 5A, second row.

**Figure 5.**
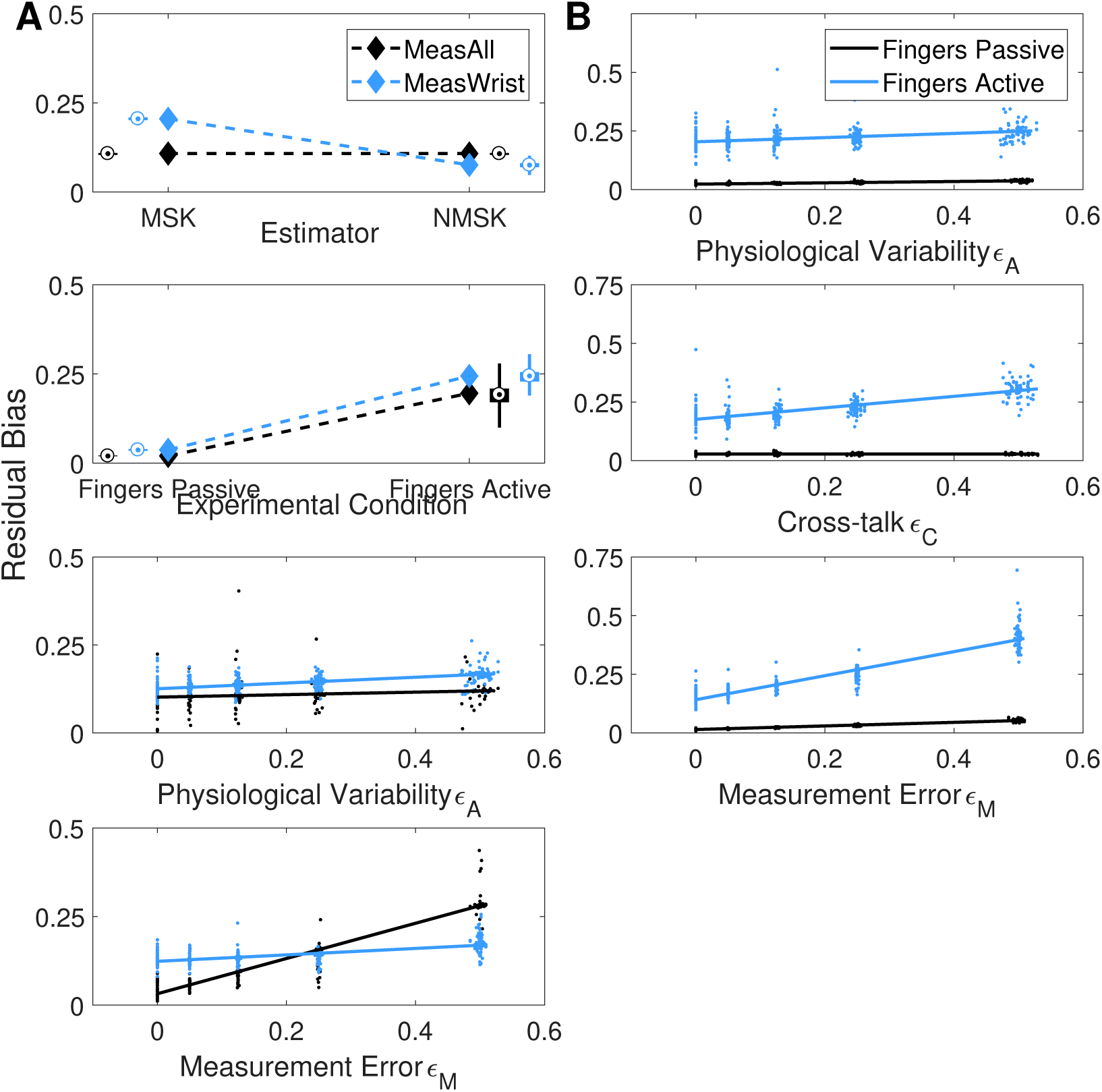
Graphical representation of the statistically significant two-way interaction terms. In the top two rows the diamonds and the lines represents the model estimates, while the simulated data reported in term of error bars or scattered dots. All plots are obtained in the average condition of noise for the unreported variables. In the columns A and B there are reported the statistically significant two-interaction terms that involve the factors Me_1_ and Ex, respectively.

For the interaction between Me_1_, Ex, and Me_3_ (Fig. 6, bottom row), measurement error was significantly associated with bias in all cases when fingers were active (0.96±0.11 MeasAll, and 0.15±0.09 MeasWrist), while a significant association was measured only in the MeasAll condition when fingers were passive (slope: 0.1±0.08 MeasAll, 0.5±0.09 MeasWrist). The association between measurement error and bias was significantly modulated by factor Me_1_: in the MeasAll condition, the slope was greater than in the MeastWrist condition (p < 0:05).

**Figure 6.**
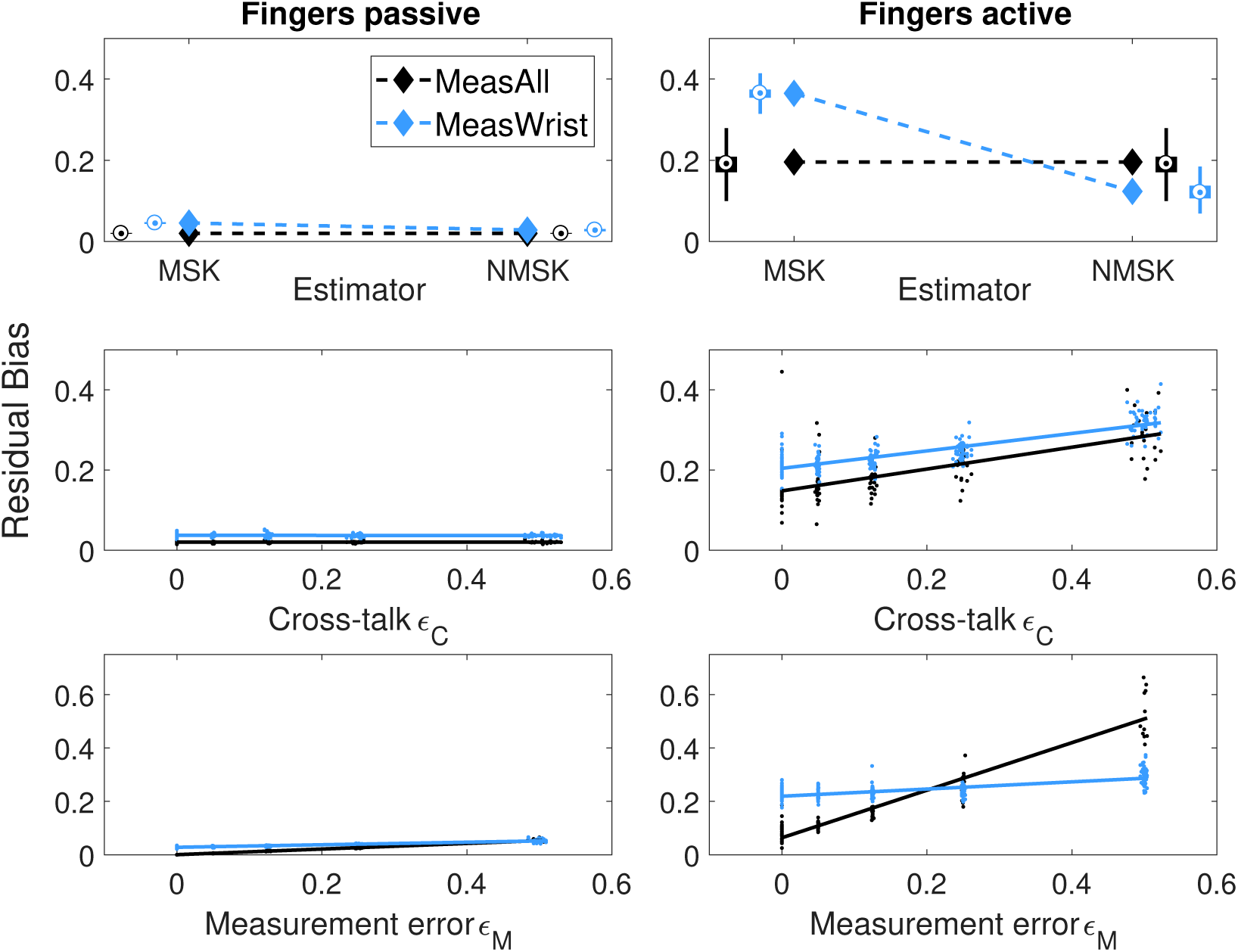
Graphical representation of the statistically significant three-way interaction terms. In the top row the diamonds represents the model estimates, with the simulated data reported in the error bar. In the central and bottom rows the lines represents the model estimates while the points are the simulated data. All plots are obtained in the average condition of noise for Ph, Me_2_ and Me_3_

Lastly, for the three four-way interactions (Fig. 7), estimation bias obtained with the NMSK estimator was not significantly different from the one obtained with the MSK estimator in the absence of finger muscles, and in the MeasAll condition when fingers are present. This was expected because of the presence of a significant effect of E only in the fingers active, MeasWrist condition. In this condition, further breakdown of the three-way interaction by variability/noise source showed that the three-way interaction is significant because of a different effect of Me_3_, whereby the slope between measurement error and bias is greater in the NMSK estimator than in the MSK estimator, in the finger active, MeasWrist condition (slope: 0.32±0.13 NMSK vs. −0.03±0.13 MSK). For physiological variability, instead, the slope between these variables and bias did not change significantly between estimators in the finger active, MeasWrist condition (slope: 0.10±0.12 NMSK vs. 0.12±0.12 MSK). Most notably, the effect of Ph on bias was not significant for neither estimator at the selected significance level. For cross-talk, instead, a signifiant association with bias was observed for both estimators (slope: 0.25±0.12 NMSK vs. 0.22±0.1 MSK), while no significant difference in slope between estimators was detected at the selected significance level.

**Figure 7.**
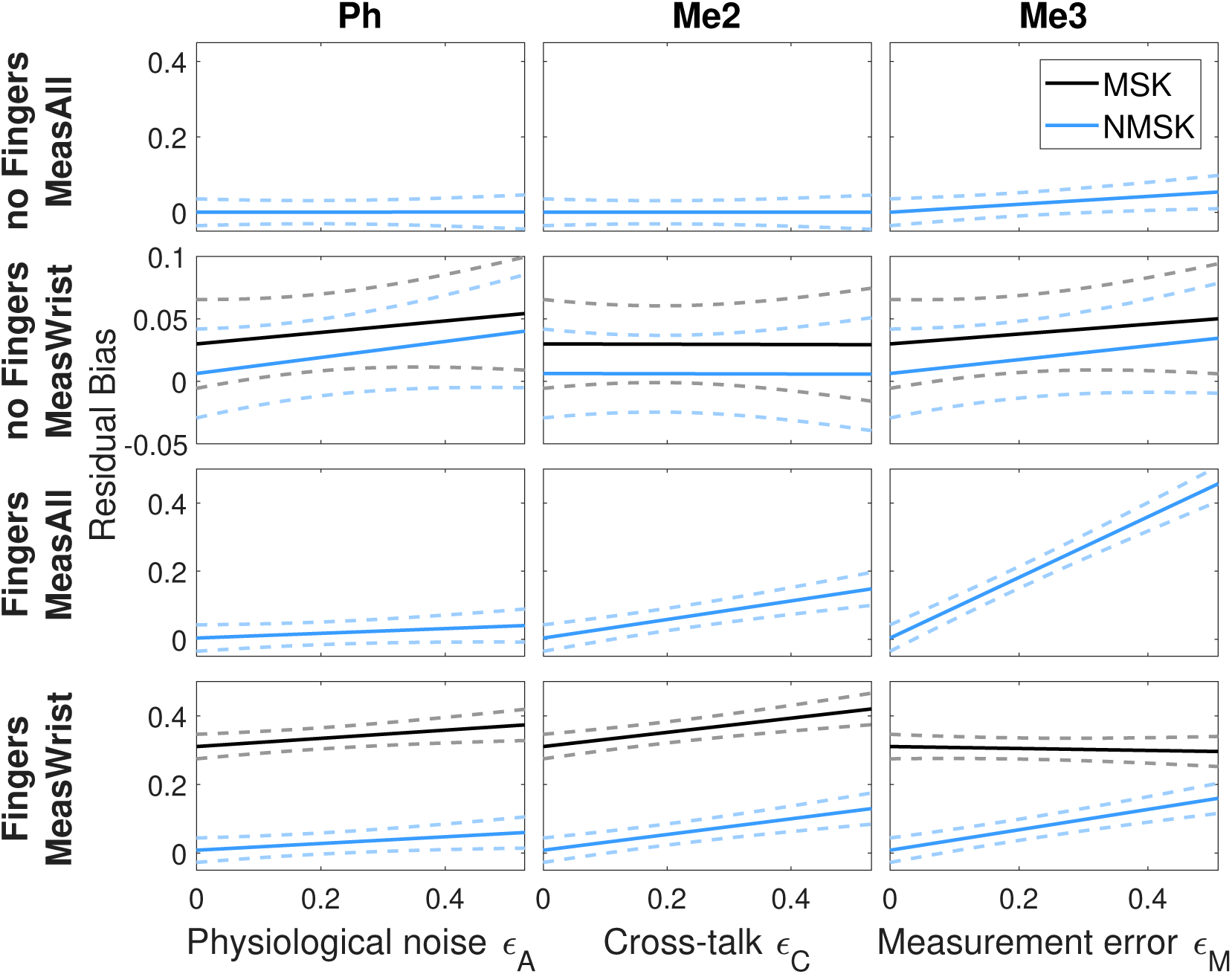
Graphical representation of the four-way interaction terms between the three categorical variables (E, Ex, Me_1_) and Ph (first column), Me_2_ (second column), and Me_3_ (third column). In each plot the sold line represents the model estimate and the dashed lines the 95% confidence intervals.

## 4. Discussion

In this study, we proposed a novel neuromusculoskeletal estimator (NMSK) that combines a standard forward dynamics estimation approach with a neural model that determines the activation of unmeasured muscles, using an optimization-based redundancy solver. We hypothesized that this novel estimator would be able to estimate individual muscle force with greater accuracy compared to a standard musculoskeletal estimator (MSK), when measurements of muscle activity are not available for all muscles.

To test our hypothesis, we ran a sensitivity analysis to quantify the effect of a set of experimental, physiological, and measurement conditions on the accuracy of muscle force estimation, both for our NMSK estimator and for the MSK estimator. To this goal, we developed a computational framework capable of simulating virtual measurements of muscle activity under a variety of experimental conditions, physiological variability and measurement error. Then, by fitting the estimation error to a general linear model, we established the association between different experimental factors and the estimation bias that characterize each estimator.

Overall, our statistical analysis demonstrated that the MSK estimator, used in standard procedures to study control of the wrist degrees of freedom (Buchanan et al. 2005; Mogk and Keir 2003), produces biased estimates of individual muscle force in presence of activity of finger muscles. However, when measurements are available from only the five main wrist muscles, the estimation bias obtained with the NMSK estimator is significantly smaller than the one obtained with the MSK estimator. Our statistical analysis further demonstrated that the effect of cross-talk on bias is greater when fingers are active compared to when fingers are passive, for either estimator. Moreover, measurement error has a greater effect on bias when fingers are active compared to when fingers are passive, but the effect is greater when measurements are obtained from all muscles. Finally, while our NMSK estimator is heavily model-based, the lack of a significantly different association between physiological variability and bias (in all conditions) provides an indication of the robustness of the NMSK estimator with respect of violations of its assumptions in the identification of the optimal solution to muscle redundancy. A more detailed discussion on our main findings is reported below.

### 4.1. Effect of estimator

Our analysis did not detect a main effect of factor estimator (E). In fact, estimator type has a significant effect on bias only under one condition: specifically when measurements are taken only from wrist muscles (MeasWrist) (Fig 6). This was expected, because when measurements are available from all muscles (MeasAll), the MSK and NMSK estimators are identical. In MeasWrist, the NMSK estimator generally reduces estimation bias over the MSK estimator (change in bias: 2±1% for finger passive, 24±3%for fingers active in the average noise conditions). The greater effect seen in the active condition is justified by the fact that in the fingers active condition, the set of unmeasured muscles is composed by n =10 muscles, while in the passive condition there is only one unmeasured muscle (i.e. PL).

A second effect of factor estimator was observed in the difference in slope of the relationship between measurement error and bias measured in the finger active, MeasWrist condition for the two estimators (slope: 32±13% NMSK vs −3±13% MSK). While this result indicates that the NMSK estimator is more sensitive to measurement error than the MSK estimator, the bias obtained using the NMSK estimator is smaller than the one obtained using the MSK estimator for all values of measurement error in the considered range.

Notably, no interaction between estimator and physiological variability was detected. As such, physiological variability is not significantly associated with bias for either estimator (slope: 10%±12%for NMSK vs. MSK 12%±12%). While this effect was expected for the MSK estimator, this was unexpected for the NMSK since this estimator is based on a neural model of muscle activation, and physiological variability effectively quantifies the distance between the assumed model and experimental conditions. The lack of a significant difference between the slopes associated with the two estimators indicates that the NMSK estimator does not provide a worse estimation compared to the standard MSK estimator even under strong violations of the validity of the assumed neural model. This observation provides an indication of the robustness of the NMSK estimator to changes in muscle co-contraction strategy. This observation is justified by the fact that also the assumption under the MSK estimator is an implicit neural model, i.e. that all muscles that are not measurable are inactive. This is a possibly inefficient solution to the muscle redundancy problem. Our analysis demonstrates that a model-based estimator that is based on a largely inaccurate neural model outperforms the standard MSK estimator based on such an implicit neural model.

### 4.2. Effect of number of measured muscles

Our analysis detected a main effect of the number of measured muscles (Me_1_). Unsurprisingly, estimation performed using the complete set of muscles yielded a smaller bias compared to the one performed based on a restricted set of muscles (bias: 11%±1 MeasAll vs. 14%±1 for MeasWrist).

Factor Me_1_ showed a significant interaction with cross-talk and experimental condition (Fig. 6, middle row). Specifically, the interaction showed that bias is associated with cross-talk only if the finger muscles are in the active condition. While this result certainly depends on the choice of the cross-correlation matrix C, it confirms previous analyses (Mogk and Keir 2003; Kong et al. 2010) that suggested how the presence of finger muscles could introduce significant cross-talk in sEMG calibration studies from the forearm.

Factor Me_1_ showed a significant interaction with measurement error and experimental condition (Fig. 6, bottom row). Specifically, the interaction showed that bias is only associated with measurement error if the finger muscles are active. Interestingly, in this condition model fit lines intersect, indicating that if measurement error is high, bias is smaller if measurements are available from a reduced set of muscles. This effect is attributable to the fact that in high measurement error conditions, half of the estimated data points are extracted using the NMSK estimator, which integrates the noisy measurements with model-based estimates, which lead to smaller estimation error (Fig. 7 - bottom right). In the MeasAll condition, instead, those same measurements are available with higher error, which leads to greater estimation error.

## 5. Conclusion

Our analysis shows that estimating the forces applied by the forearm muscles during isometric tasks that involve the wrist joint can be challenging. In fact, by relying only on the EMG measurements obtained from the five main wrist muscles, when fingers are active (i.e. MSK estimator), highly biased estimates of individual muscle force are obtained (average bias of 36±2%). Even though leaving the fingers unconstrained decreases estimation bias, this procedure is not always practically possible and it still requires to train subjects to not activate finger muscles during data collection. Instead, the integration in the estimator of a neural model of the optimal muscle co-contraction strategy (i.e. NMSK estimator) reduces the estimation bias by about 25% of the true value. Moreover, even when the validity of the neural model used for the NMSK estimator is compromised, the NMSK estimator still outperforms standard estimators based on measurements of a partial set of muscles.

